# Loss of α-actinin-3 during human evolution provides superior cold resilience and muscle heat generation

**DOI:** 10.1101/2020.10.03.323964

**Authors:** VL Wyckelsma, T Venckunas, PJ Houweling, M Schlittler, VM Lauschke, CF Tiong, H Wood, N Ivarsson, H Paulauskas, N Eimantas, DC Andersson, KN North, M Brazaitis, H Westerblad

## Abstract

The fast skeletal muscle protein α-actinin-3 is absent in 1.5 billion people worldwide due to homozygosity for a nonsense polymorphism in the *ACTN3* gene (R577X) ^1^. The prevalence of the 577X allele increased as modern humans moved to colder climates, suggesting a link between α-actinin-3 deficiency and improved cold tolerance ^1,2^. Here, we show that humans lacking α-actinin-3 (XX) are superior in maintaining core body temperature during cold-water immersion due to changes in skeletal muscle thermogenesis. Muscles of XX individuals displayed a shift towards more slow-twitch isoforms of myosin heavy chain (MyHC) and sarcoplasmic reticulum (SR) proteins, accompanied by altered neuronal muscle activation resulting in increased tone rather than overt shivering ^3,4^. Experiments on *Actn3* knockout mice showed no alterations in brown adipose tissue (BAT) properties that could explain the improved cold tolerance in XX individuals. Thus, this study provides a clear mechanism for the positive selection of the *ACTN3* X-allele in cold climates and supports a key thermogenic role of skeletal muscle during cold exposure in humans.

## Main

The sarcomeric protein α-actinin-3 resides in the Z-discs of fast skeletal muscle fibers, where it cross-links the actin filaments of adjacent sarcomeres ^5–7^. The lack of functioning *ACTN3* does not cause muscle disease, but it has been shown to affect muscle function both in the general population and in athletes ^8,9^; in general, α-actinin-3 deficiency is detrimental for power and sprint activities ^8,10,11^.

A study exploring evolutionary implications of α-actinin-3 deficiency demonstrated that the X-allele became more abundant as humans migrated out of Africa into the colder climates of central and northern Europe ^1,2,12^. This led to the hypothesis that α-actinin-3 deficient humans are superior in adapting to lower temperature ^2^. On this basis, we sought to determine whether α-actinin-3 deficient (XX) humans were better at defending their body temperature during an acute cold challenge than humans with functioning *ACTN3* (RR). Young male XX and RR individuals (Supplementary Table 1) were immersed in 14 °C water for 20 min periods interposed by 10 min pauses in room-tempartured air; cold-water exposure was continued until the rectal temperature reached 35.5 °C or for a total of 120 min (170 min including the pauses). The percentage of individuals able to maintain their body temperature above 35.5 °C degrees for the complete cold-water exposure was markedly higher in the XX group (69%) than in the RR group (30%) (Fig. 1A). The average rate of decline of rectal (T_re_) and gastrocnemius muscle (T_mu_) temperatures in XX subjects was about half of that in RR subjects (Fig. 1B and 1C). The rate of skin temperature (T_sk_) decline, on the other hand, was not significantly different between the two groups (Fig. 1D). The overall increase in energy consumption induced by the cold challenge was assessed by measurements of heart rate and the rate of respiratory O_2_ uptake (VO_2_) and CO_2_ exhalation (VCO_2_); all three rates were significantly increased at the end of cold exposure irrespective of ACTN3 genotype (Fig. 1E-G). Thus, α-actinin-3 deficient individuals showed superior protection of core body temperature during an acute cold stress and this was achieved without increased energy consumption as judged from similar cold-induced increases in heart rate, VO_2_ and VCO_2_ in XX and RR individuals.

**Fig. 1.**
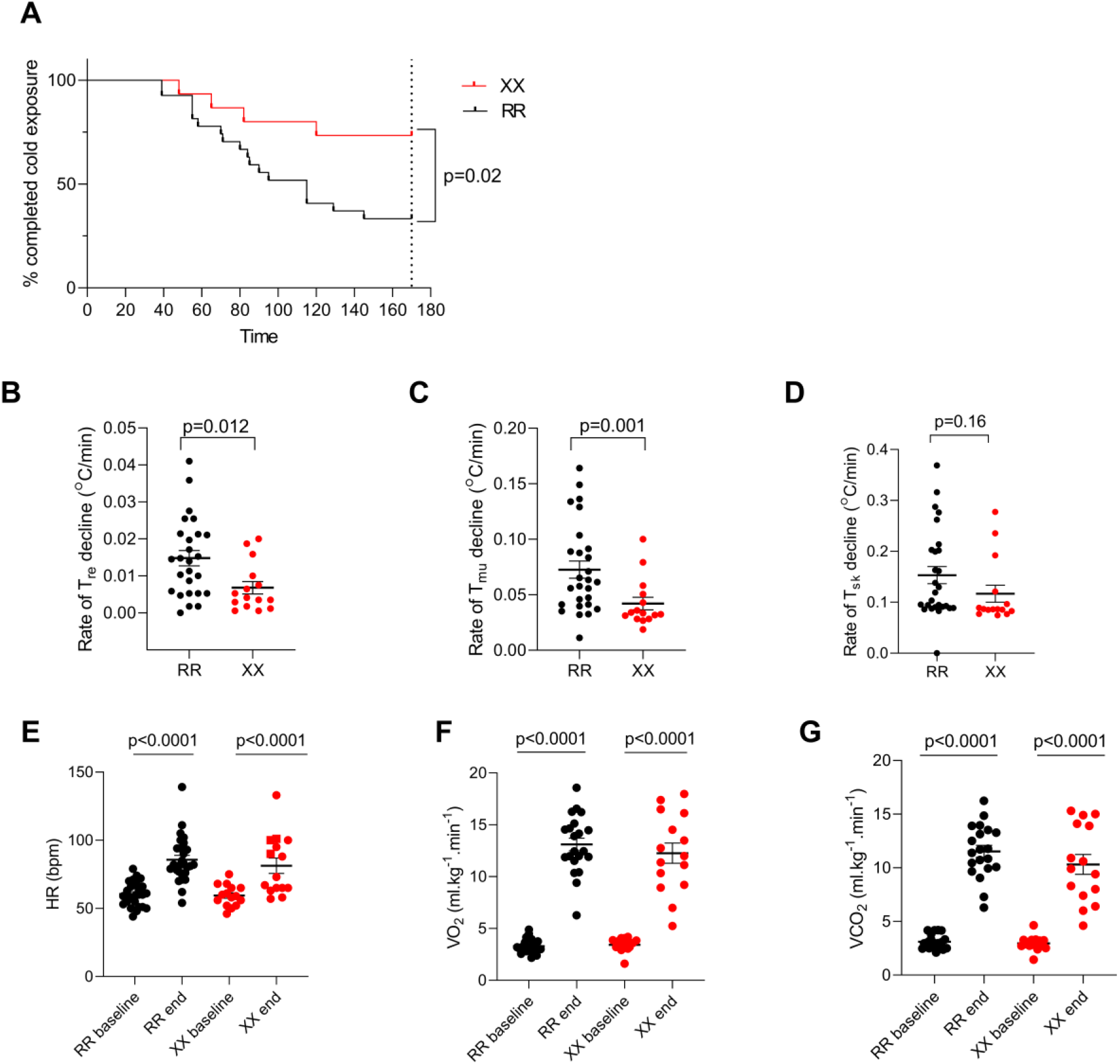
Temperature measurements and physiological responses during cold-water immersion. **A)** Survival plot of the time taken to reach a rectal temperature (T_re_) of 35.5 °C or sustaining the complete 170 min period of cold-water immersion in RR (n = 27) and XX (n = 15) individuals. Log-rank (Mantel-COX) test was used to assess statistical difference between RR and XX individuals. The decline rate in rectal (T_re_; **B**), intramuscular (T_mu_; **C**) and skin (T_sk_; **D**) temperatures in RR and XX subjects. Statistical difference between the two groups was assessed with unpaired t-test. Heart rate (**E**) and rate of pulmonary O_2_ uptake (VO_2_, **F**) and CO_2_ exhalation (VCO_2_, **G**) before (baseline) and at the end of cold-water immersion in RR and XX subjects. Statistical assessment with 2-way RM ANOVA revealed no differences between the two groups either before or at the end of cold-water exposure. Plots show values for each RR (black circles) and XX (red circles) individual and mean ± SEM.

In mice, *Actn3* knockout (KO) results in changes in intracellular Ca^2+^ handling with marked increases in SR Ca^2+^ leak and the subsequent heat-generating active SR Ca^2+^ re-uptake via the SR Ca^2+^-ATPase (SERCA) ^13^. Uncoupling of SERCA activity from the actual Ca^2+^ transport into the SR is considered a key component in muscular non-shivering thermogenesis ^14–17^. SERCA is expressed in several different isoforms in mammalian tissues with SERCA1 and SERCA2a being the main isoforms in adult fast-twitch and slow-twitch muscle fibers, respectively ^18–20^. Notably, we detected a shift in dominance from SERCA1 in RR muscles to SERCA2a in XX muscles (Fig. 2A-B). The SR Ca^2+^ storage protein, calsequestrin (CSQ), also shows a fiber type-dependent isoform distribution with CSQ1 dominating in fast-twitch fibers and CSQ2 in slow-twitch fibers ^19,21^. We observed similar total CSQ expression in XX and RR muscles, whereas the expression of CSQ2 was about twice as high in XX compared to RR muscles (Fig. 2C-D). The difference in SERCA and CSQ isoform expression between XX and RR muscles may reflect a larger volume of the muscle to be composed of slow-twitch fibers in XX than in RR subjects. Therefore, we used high-sensitivity silver staining to analyze the MyHC composition and found that XX muscles had significantly more slow MyHC I (β-MyHC) and less fast MyHC IIx than RR muscles (Fig. 2E).

**Fig. 2.**
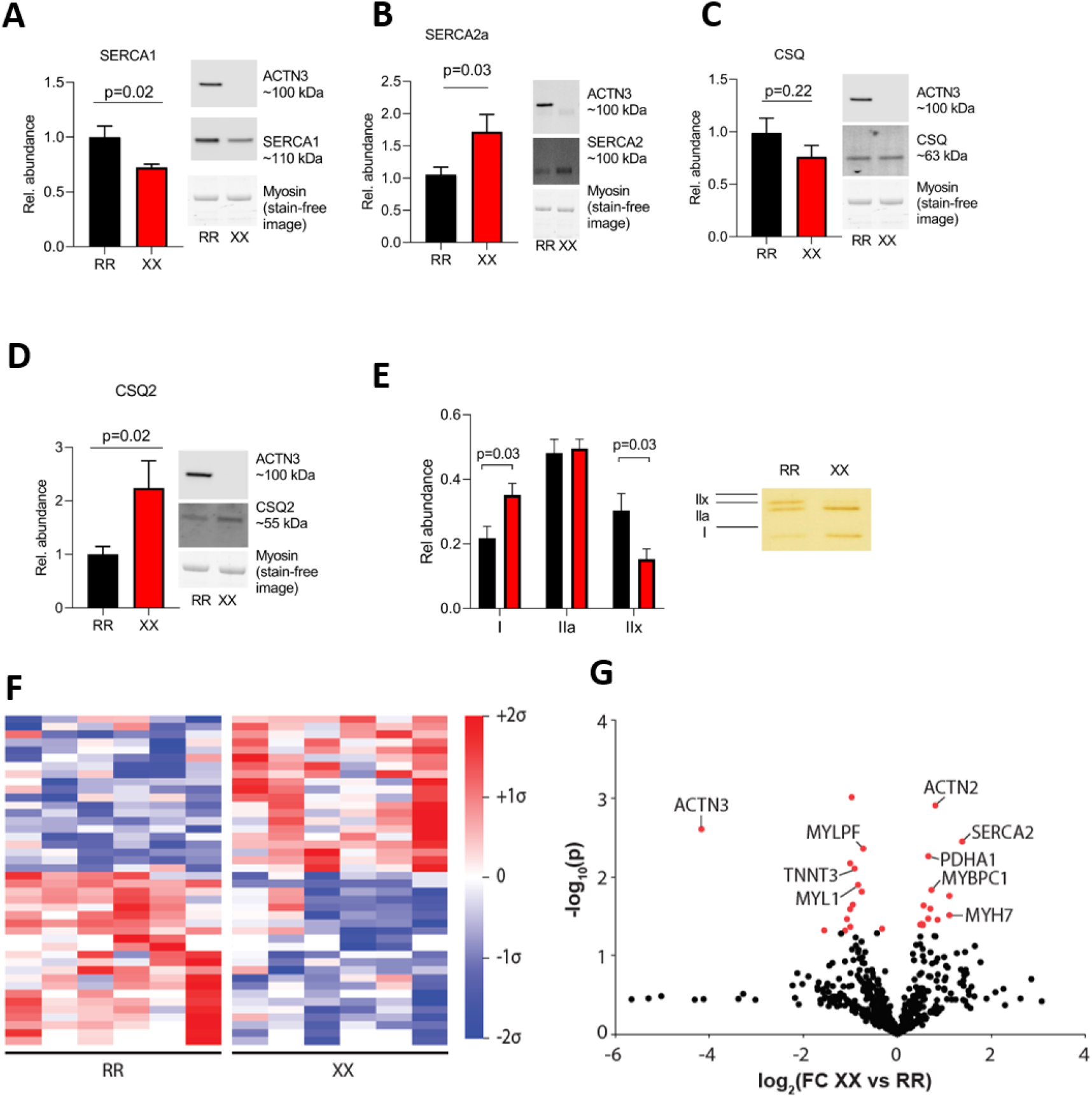
ACTN3 deficiency is accompanied by a shift towards a slower skeletal muscle phenotype. Summary data (mean ± SEM) and representative western blots of the SR Ca^2+^-handling proteins SERCA1 (**A**), SERCA2a (**B**), CSQ (**C**), and CSQ2 (**D**) in muscle of RR (n = 8) and XX (n = 7) individuals. Band intensities were normalized to their respective myosin loading controls. Data expressed relative to the mean value of the RR group, which was set to 1.0. Statistical difference between the two groups was assessed with unpaired t-test. **E)** Silver stained gels were used to assess the distribution of MyHC isoforms in RR (n = 7) and XX (n = 7) individuals. Right part shows a representative example of the distribution of MyHC in RR and XX individuals. The total staining of the three MyHC bands was set to 1 in each subject. Statistical difference between the two groups was tested with unpaired t-test. **F)** Mean-centered sigma-normalized heatmap of differentially expressed proteins (P < 0.05). **G)** Volcano plot of all identified proteins (n = 601) expressed as fold-change (FC) in XX compared to RR individuals. Differentially abundant proteins are indicated in red.

The sarcomeric α-actinins are known to interact with a multitude of functionally diverse proteins involved in structural, metabolic, signaling, and Ca^2+^-handling pathways ^10^. We used proteomics as an exploratory measure to look for further differences in muscle between XX and RR individuals. This proteomic analysis of skeletal muscle biopsies shows distinct differences in protein abundance between XX and RR individuals. Overall, we found 42 proteins to be differentially expressed between genotypes (Fig. 2F), including higher protein levels of slow-twitch muscle fiber markers (MyHC I (MYH7), slow-type myosin-binding protein C (MYBPC1), SERCA2) and lower levels of fast-twitch markers (myosin light chain 1/3 (MYL1), myosin regulatory light chain 2 (MYLPF), fast-type muscle troponin T (TNNT3)) in XX than in RR individuals (Fig. 2G). Furthermore, pathway analysis revealed that differentially expressed proteins were enriched in pyruvate metabolism (*P* = 0.0003; FDR = 0.02), including pyruvate dehydrogenase (PDHA1) with an ~60% higher protein expression in XX than in RR individuals.

Although the difference in SERCA and CSQ isoform expression between XX and RR muscle homogenates corresponds with the MyHC distributions, α-actinin-3 deficiency might still affect the abundance of these Ca^2+^-handling proteins within individual fiber types. However, fiber type-specific western blots performed on pooled single muscle fibers did not reveal any fiber type-dependent differences in the expression of SERCA1, total CSQ or CSQ2 between XX and RR muscles (Supplementary Fig. 1). Intriguingly, SERCA2a was higher in the RR than in XX Type I fibers, which might indicate that the abundance of SERCAs is not the limiting factor in temperature regulation. SERCA might be involved in muscular non-shivering thermogenesis via its associated protein sarcolipin (SLN), which interferes with SERCA function by mediating uncoupling of the SR Ca^2+^ uptake from the heat-generating ATP hydrolysis ^15,22^. Hence, we measured SLN expression in muscle homogenates and in pooled single fibers, and the results showed no difference in SLN expression between XX and RR muscles (Supplementary Fig. 2 and Supplementary Fig. 3). Another potential Ca^2+^-SR-related mechanism for heat generation is Ca^2+^ leak through the SR Ca^2+^ release channel, the ryanodine receptor 1 (RyR1) channel complex, due to dissociation of the channel-stabilizing subunit FK506 binding protein (FKBP12) ^23,24^. However, immunoprecipitation experiments did not reveal any difference in the amount of FKBP12 bound to RyR1 between XX and RR muscles (Supplementary Fig. 2B). To summarize, we could not detect any SR Ca^2+^-handling protein-dependent explanation for the superior cold tolerance of XX subjects.

Brown adipose tissue is an important heat-producing thermo-effector in mammals ^25^. However, adult humans have relatively little brown adipose tissue (BAT) and it is therefore difficult to assess its role as a heat generator ^26^. We utilized the well-defined *Actn3* KO mouse model to examine the impact of BAT activation as a mechanism for improved heat generation in α-actinin-3 deficient individuals ^1^. Wildtype (WT) and *Actn3* KO mice were kept in a cold (4 °C) room for 5 hours and core body temperature was measured at regular intervals using a rectal probe. In accordance with the human results, the number of mice able to maintain their body temperature above 35.5°C for the cold exposure period was markedly higher in the *Actn3* KO (41%) than in the WT (16%) group (Fig. 3A). Although the overall rate of temperature decline during cold exposure was not significantly different between the two groups (Fig. 3B), *Actn3* KO mice were significantly lighter than their WT counterparts (Fig. 3C) and therefore showed an improved cold tolerance after normalizing for body weight (Fig. 3D). Following cold exposure, *Actn3* mRNA was present in BAT of WT mice, but absent in the KO mice (Fig. 3E). RNA-sequencing was performed on BAT collected from mice subjected to either thermal neutrality (TN, 30 °C), room temperature (RT, 22 °C) or low temperature (cold, 4 °C). An unbiased principal component analysis (PCA) and heat map clustering separated samples based on temperature (TN, RT and cold), whereas it showed no effect of *Actn3* genotype (Fig. 3F and G). Further analysis showed over 2000 differentially expressed genes in BAT following cold exposure (Fig. 3H); however, only one transcript was significantly altered based on genotype and temperature (cold; glycine/arginine rich protein 1; Grrp1), excluding physiologically relevant genotype-specific differences in BAT properties following acute cold exposure (Fig. 3I). To sum up, these data imply that the improved cold tolerance observed in α-actinin-3 deficient humans and mice is due to improved skeletal muscle heat generation without any detectable influence of BAT.

**Figure 3:**
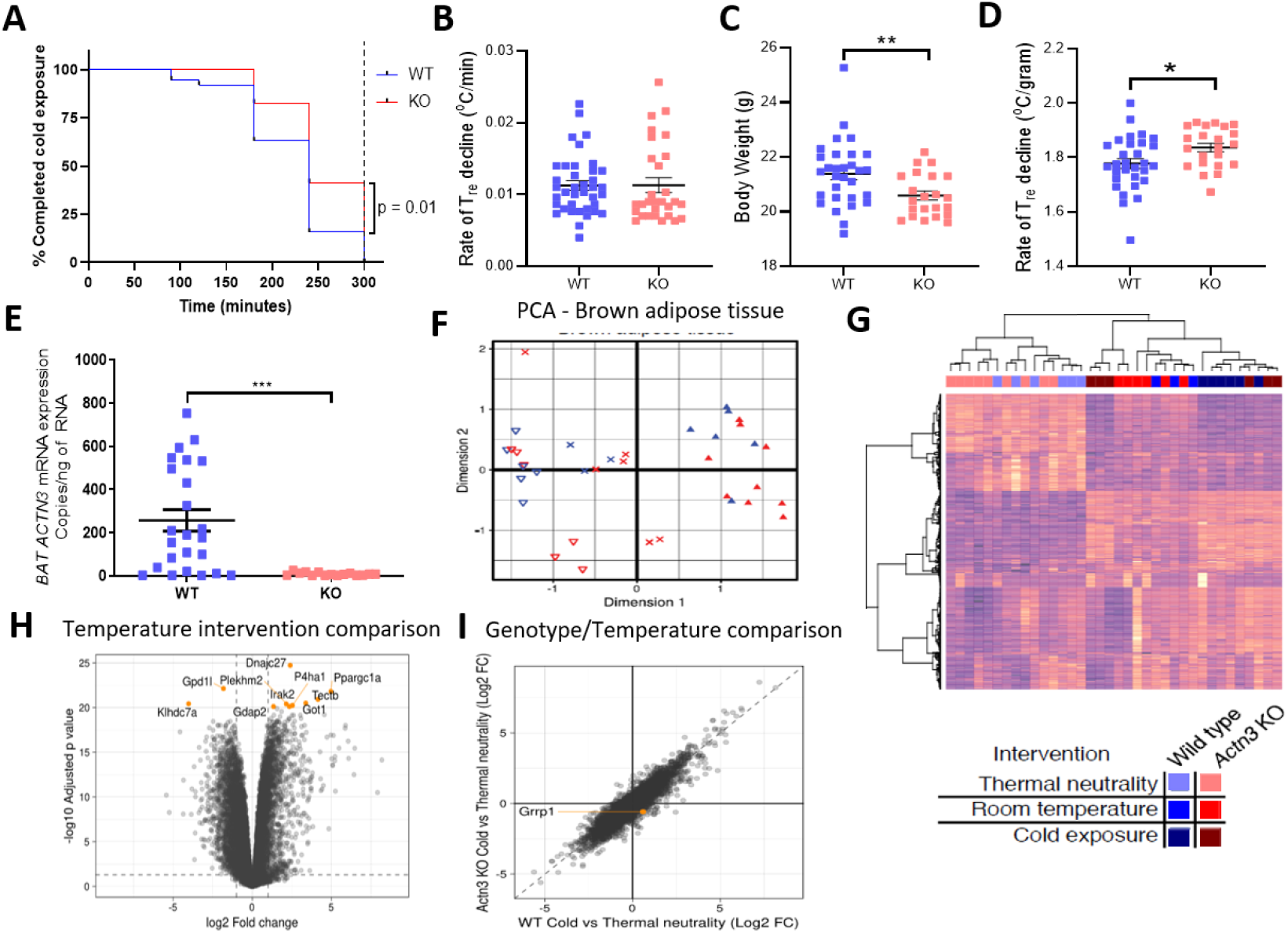
Improved cold tolerance in *Actn3* KO mice not related to altered BAT properties. **A**) Survival plot of the time taken to reach 35.5°C core body temperature over a 5 hours period exposed to 4 °C air temperature in WT (n = 38) and Actn3 KO (n = 29) mice. Log-rank (Mantel-COX) test was used to assess statistical difference between WT and KO mice. The overall rate of decline in rectal body temperature (**B**), body weight (**C**) and rate of rectal body temperature decline per g body weight (**D**). **E**) Following cold exposure, Actn3 mRNA is present in WT but not in Actn3 KO mice. RNA-sequencing analyses show an effect of temperature but no effect of Actn3 genotype on BAT activation with both the principal component analysis (PCA, **F**) and heat map (**G**). **H**) Volcano plot of altered genes confirms that marked changes in BAT gene expression occurs following acute cold exposure, with >2000 differentially expressed genes identified. **I**) Interaction plot shows no differences based on Actn3 genotype in BAT following acute cold exposure. *P < 0.05, **P < 0.01, ***P < 0.001 with unpaired t-test.

The major heat-generating mechanism in mammalian skeletal muscle during an acute cold challenge is involuntary activation of motor units resulting in skeletal muscle contraction. This mechanism is generally referred to as shivering thermogenesis although it involves both increased basal muscle tone due to continuous low-intensity activation and overt shivering due to high-intensity bursting activity ^3^. In our human cohort, we used surface electromyography (EMG) to follow the activation of pectoralis major muscles during cold-water immersion and observed more frequent bursting activity in RR individuals with mean data showing an approximately two times higher rate of bursts in RR than in XX muscles (Fig. 4A and B); the markedly higher activity during bursts also resulted in a slightly higher mean EMG signal frequency in RR muscles (Fig. 4C). On the other hand, there was no significant difference in the amplitude of the EMG signal, which reflects the overall number of muscle fibers being activated (Fig. 4D). Interestingly, cold-induced low-intensity continuous muscle activity is associated with activation of type I muscle fibers, whereas high-intensity burst activity is linked to recruitment of type II muscle fibers ^4,27,28^. Thus, the difference in burst rate between XX and RR muscles is consistent with the difference in fiber type distribution with more type I and less type II MyHC in XX than in RR muscles (see Fig. 2E).

**Figure 4.**
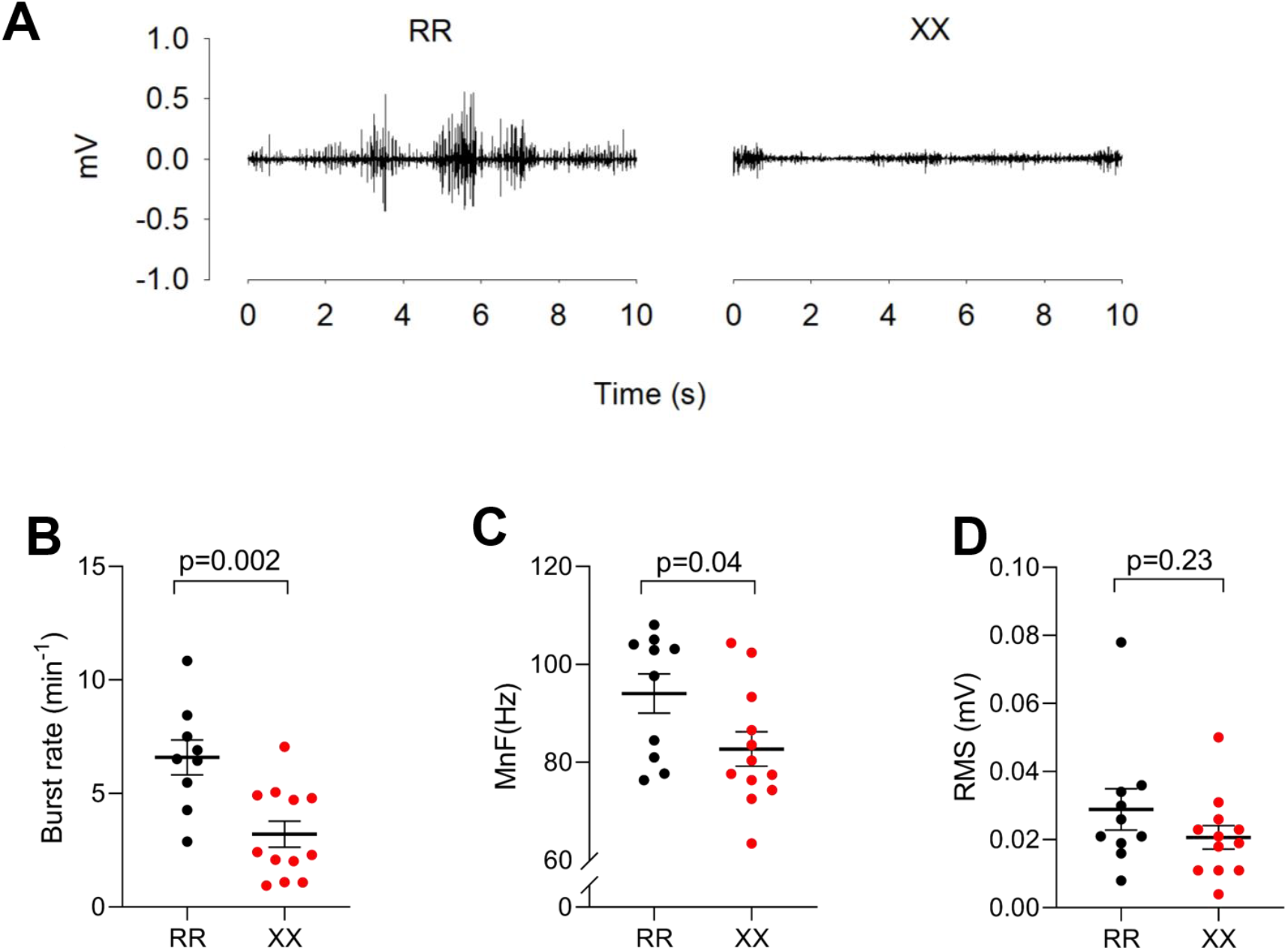
Bursting muscle activity rather than increased muscle tone is more prominent in RR muscles. **A)** Representative EMG records from pectoralis major muscles during cold exposure showing continuous low-intensity activity in the XX individual and frequent bursts of high-intensity activity in the RR individual. Summary data of the burst rate (**B**), mean EMG signal frequency (MnF; **C**), and amplitude (RMS, root mean square; **D**). Plots show values for each RR (black circles) and XX (red circles) individual and mean ± SEM. Statistical differences between RR and XX individuals were assessed with unpaired t-test.

In conclusion, α-actinin-3 deficient humans show improved cold tolerance during cold water immersion, which is associated with a shift towards more slow-twitch type I MyHC. During cold exposure, motor neuron activation occurs mainly as an effective heat-generating increase in muscle tone rather than overt shivering ^4^. During prolonged cold exposure, the improved heat generation via increased muscle tone suggests that XX individuals would consume less energy and be less susceptible to developing muscle fatigue compared to individuals that express α-actinin-3 (RR), providing an evolutionary survival advantage for XX individuals. This mechanism provides an explanation for the increase in X-allele frequency as modern humans migrated from Africa to the colder colder Euroasian climate over 50.00 years ago ^2,12^. However, cold tolerance is seldom a key issue in modern societies and an energy efficient phenotype can be problematic in the context of a current lifestyle with high caloric intake and reduced physical activity.

## Acknowledgments

We thank Monika Kisieliute and Andreius Subocius for their assistance in genotyping and collection of human muscle biopsies. We also thank Sophie Agius and Alison Burns for their contribution to maintaining the *Actn3* KO mouse colony.

## Author Contribution

TV, MB, HWe conceived the human study. VLW, TV, MS, NI, HP, NE, MB performed experiments in the human study. VLW, TV, MS, VL, NI, HP, NE, DCA, MB, HWe analysed and intrepreted human data. KNN conceived the *Actn3* KO mouse study. PJH, CFT, HWo completed the mouse analyses and RNA-sequencing. VLW, PJH, KNN, HWe drafted the manuscript. All authors provided critical evaulation of the manuscript and approved the final version.

## METHODS

### Participants, mice and ethical approval

#### Humans

Healthy young (18-40 yrs) males were recruited to participate in the study. Before being included in the study, each participant was informed of the aims, the experimental procedures and the potential risks of the study and signed a written informed consent form consistent with the principles outlined in the Declaration of Helsinki. The study was approved by Kaunas Regional Biomedical Research Ethics Committee (license number BE-2-30).

The participants were moderately physically active (< 2 h physical exercise / week) and did not participate in any formal physical exercise or sport program. They had not been involved in any temperature manipulation programme or extreme-temperature exposure for at least 3 months. Individuals with any existing medical condition or taking medication that could affect natural thermoregulation were excluded from the study. The physical characteristics of the participants are presented in **Supplementary Table 1**. Their weight (in kg), body fat percentage (TBF-300 body composition analyser, Tanita, UK Ltd., West Drayton, UK) and height (in cm) were measured, and body mass index was calculated. Body surface area (in m^2^) was estimated as previously described using the following formula: body surface are = 128.1×Weight^0.44^×Height^0.60^ ^29^. Skinfold thickness (in mm) was measured with a skinfold calliper (SH5020, Saehan, Masan, Korea) at 10 sites (chin, subscapular, chest, side, suprailium, abdomen, triceps, thigh, knee and calf) and the mean subcutaneous fat layer thickness was calculated ^30^.

#### *Actn3* KO and WT mice

All animal work was carried out in accordance with approval from the Murdoch Children’s Research Institute Animal Care and Ethics Committee (Approval No. A760). Animals were housed in a specific pathogen-free environment at a constant ambient temperature of 22 °C and 50% humidity on a 12 h light-dark cycle, with *ad libitum* access to food and water, unless otherwise specified. Generation of *Actn3* KO mice on a C57BL/6J background has been previously reported ^31^. Age-matched female WT and *Actn3* KO littermates derived from heterozygous *Actn3* crosses were used for all animal studies.

### Genotyping

#### Human

DNA was extracted from blood samples using the NucleoSpin Blood kit (Macherey-Nagel, GmbH & Co. KG, Düren, Germany) according to the manufacturer’s protocol. *ACTN3* R577X genotype was determined using a PCR-RFLP method as previously described^32^.

#### Mouse

*Actn3* WT and KO genotypes were determined using PCR-RFLP using DNA extracted from ear punch biopsies and extracted as outlined previously ^1^.

### Acute cold exposure

#### Human cold-water immersion protocol

An intermittent whole-body water immersion cooling protocol was used as previously described ^33–35^. All experiments were conducted indoors at the same time of day (from 7:00 a.m. to 11:00 a.m.). The participants refrained from consuming any food for at least 12 h before the experiment. To standardize the state of hydration and the sensation of thirst, subjects were allowed to drink still water as desired until 60 min before the water-immersion session. The experiments were performed at a room temperature of 22 °C and a relative humidity of 60%. Prior to cold-water immersion, the participants rested for 10–15 min dressed in a T-shirt, swim shorts and socks and baseline ventilation parameters, heart rate, and temperatures were measured during the subsequent 20 min. Thereafter, they entered a 14 °C water bath with only the head above the surface. Individuals stepped out of the bath every 20 min and rested for 10 min at room temperature, and then returned to the water bath for the next 20 min of cold-water immersion. This intermittent whole-body water immersion procedure continued until either the rectal temperature (T_re_) had decreased to 35.5 °C or a maximum of 120 min of cold-water immersion (170 min including the breaks).

#### Mouse acute thermoneutral and cold exposure

For acute temperature exposure experiments, 12 week old female WT and *Actn3* KO mice were singly housed in cages kept at either 30 °C or 4 °C as previously published ^36^. Briefly, mice housed at thermoneutrality (30 °C) were acclimatised at this temperature for 20 h (with food and water *ad libitum*) prior to commencement of experiments. Food, water and bedding were removed from cold-exposed mice during the 5 h cold exposure period.

### Body temperature measurements

#### Humans

T_re_ was measured throughout the experiment using a thermocouple (Rectal Probe, Ellab, Hvidovre, Denmark; accuracy ± 0.1 °C) which was inserted by the subjects to a depth of 12 cm past the anal sphincter. Muscle (T_mu_) and skin (T_sk_) temperatures were measured before and at the end of the water immersion session. The T_mu_ was measured with a needle microprobe (MKA, Ellab; accuracy ± 0.01 °C) inserted ~3.5 cm under the skin covering the lateral gastrocnemius muscle of the right leg. The skin was prepared before each intramuscular temperature measurement by shaving and disinfecting with a cotton-wool tampon soaked with medicinal alcohol. The insertion area was marked to ensure the repeatability of the measurement. T_sk_ was measured with thermistors taped to the back, the thigh and the forearm (DM852, Ellab; accuracy ± 0.1 °C), and mean T_sk_ was calculated as: T_sk_=0.5T_back_+0.36T_thigh_+0.14T_forearm_ ^37^.

#### Mice

Core body temperature was measured by a rectal probe (BAT-12 microprobe thermometer) over a 5 hour period between the times of 8:00 and 14:00. Temperatures were measured at 0, 30, 60, 90, 120, 180, 240, and 300 min. Body weights were recorded before and after the 5 h temperature measurement period. At the conclusion of the temperature assessment, all mice were euthanized by cervical dislocation and tissues were collected for further analysis.

### Human functional measurements

#### Spirometry and heart rate measurement

A mobile spirometry system (Oxycon Mobile, Jaeger/VIASYS Healthcare, Hoechberg, Germany) was used to measure VO_2_ and VCO_2_ on a breath-by-breath basis. Automatic calibration of the gas analyser and delay time were performed before measurements as described by the manufacturer, i.e. a calibration gas at 180 kPa (15.2% O_2_, 5.02% CO_2_, and 79.62% N_2_) was supplied to attain gain, offset, and delay times within 1%. Heart rate was measured throughout the experiment with a heart rate monitor (S-625X, Polar Electro, Kempele, Finland).

#### EMG measurement of thermoregulatory muscle activation

Heat-generating muscle activation is pronounced in the chest region and we therefore measured EMG signals in the *pectoralis major* muscle during cold-water immersion ^38,39^. After careful preparation of the skin (shaving, abrading, and cleaning with alcohol wipes) to obtain a low impedance (< 10 kOhm), a surface EMG sensor (SX230W, Biometrics Co., Ltd., Gwent, UK) with integrated bipolar Ag–AgCl electrodes (10 mm diameter, 20 mm centre-to-centre distance) and differential amplifier (gain 1000, input impedance 100 MΩ, an input noise < 5 μV, common mode rejection ratio higher than 96 dB) was placed on the right pectoralis major muscle. The ground electrode (R206, Biometrics Co., Ltd.) was positioned on the wrist of the right hand. The EMG sensor and ground electrode were connected to a portable data acquisition unit (DataLog P3X8, Biometrics Co., Ltd.) Before measurements, the channel sensitivity was set to 3 V and the excitation output to 4600 mV as recommended by the manufacturer. EMG signals were digitized and files were stored on a computer for subsequent analyses of the mean frequency (MnF, in Hz) and root mean square (RMS, in mV) using a dedicated software (Biometrics DataLOG, Gwent, UK) and manual analysis of the rate of burst activity.

### Human protein analyses

#### Muscle biopsies

Prior to cold-water immersion, biopsies from the *vastus lateralis* muscle were collected from a subgroup of RR (n = 11) and XX (n = 8) individuals. The biopsy site was cleaned with alcohol and anesthetized locally. After making a small skin cut with a scalpel tip, a biopsy needle was inserted perpendicular to the muscle fibers and biopsies were collected with and automatic biopsy device (Bard Biopsy Instrument, Bard Radiology, Covington, USA). After collection, muscle biopsies were snap frozen in liquid nitrogen and stored at −80°C until analysis. The skin cut was cleaned and closed with wound closure strips. Biopsies were not collected post cold exposure.

#### Whole muscle homogenate preparation

Whole muscle homogenate was prepared from frozen muscle biopsies for western blots, analysis of MyHC isoform composition, and immunoprecipitation (IP) experiments. Approximately 15 mg of frozen muscle was weighed and homogenized on ice (1:20 w/v) in HEPES lysis buffer (20 mM HEPES, 150 mM NaCI, 5 mM EDTA, 25 mM KF, 5% Glycerol, 1 mM Na_3_VO_4_, 0.5% Triton, pH 7.6) with Protease Inhibitor (#11836145001, Roche, 1 tablet per 50 ml). After this stage, 70 μL was diluted to 33 μg wet weight muscle μl^−1^ using 3 × SDS denaturing solution (0.125 M Tris-HCI, 10% glycerol, 4% SDS, 4 M urea, 10% 2-mercaptoethanol and 0.001% Bromophenol Blue, pH 6.8). Finally, samples were further diluted to 2.5 μg wet weight muscle μl^−1^ with 1× SDS solution (3 × SDS denaturing solution diluted 2:1 with 1 × Tris.Cl (pH 6.8)). A small amount of undiluted homogenate was taken from each RR participant to make a calibration curve included on every gel for western blotting. The remaining undiluted homogenate was used for IP.

#### Single fiber collection, fiber-typing and pooling

Approximately 10 mg of muscle was freeze dried for 24 h. Biopsies were left in a desiccator in drying pearls (Sigma) at room temperature for 60 min and then placed in −20 °C for long-term storage. Between 40-60 segments of single fibers were collected from each muscle biopsy and placed in 12 μl of 1 × SDS denaturing solution, once collected fibers were kept at room temperature for 60 min and then stored in −80 °C until dot blotting.

Each single fiber segment was fiber-typed using the dot-blotting method as previously described ^40^. Briefly, a PVDF membrane was activated in 96% ethanol for 120 s and then activated in transfer buffer containing 20% methanol for 120 s. Following activation, 1 μl from each single fiber tube was spotted on to a membrane. Once the membrane had dried, membranes were reactivated in ethanol (120 s) and transfer buffer (120 s), membranes were washed in × 1 Tris-buffered saline-Tween (TSBT) and then blocked for ~10 min in 5% blocking buffer (Bio-Rad) in 1 × TBST. The membrane was then incubated in primary antibody overnight at 4 °C with 2 h at room temperature. After washing and incubation with a secondary antibody and 1 × TBST washes the membrane was coated with chemiluminescent substrate (Clarity Max ECL substrate, Bio-Rad) and imaged on Chemidoc MP (Bio-Rad).

In this study, two membranes were prepared simultaneously from each fiber segment. These membranes were probed either for MyHC II (mouse, monoclonal IgG,A4.74, Developmental Studies Hybridoma Bank (DSHB)) or for MyHC I (mouse, monoclonal IgM, A4.840, DSHB) diluted 1:200 in blocking buffer in PBS (LiCOR Biosciences) 1:1 v/v in 1x TBST. Single fiber segments of the same fiber type from each muscle biopsy were subsequently pooled into a single tube and frozen at −80 °C until western blotting. Each pool ranged between 4-15 fibers. Only fiber segents that were identified either as MyHC II or MyHC I were used in subsequent analyses.

#### Western blotting

Proteins of whole homogenates and pools of single fiber segments were separated on either 4-15% TGX stain-free gels or for analysis of SLN, 16.5% Tris Tricine gels. TGX stain-free gels had total protein visualized prior to transfer and analyzed on Image Lab software (Image Lab 6.0, Bio-Rad) ^41^. Protein was wet transferred to PVDF or nitrocellulose membrane (SLN) for 1 h. Following transfer, tris-tricine gels were stained (Coomassie Brilliant Blue R-250, Bio-Rad) for 2 h at room temperature and de-stained (40% methanol, 10% acetic acid) for 2 × 1 h washes at room temperature and then stored overnight in MilliQ H_2_O before being visualized for myosin bands on a Chemi Doc MP (Biorad). Membranes were blocked at room temperature for 2 h using LI-COR blocking buffer with TBS (LI-COR Biosciences). After blocking, membranes were incubated in primary antibody overnight at 4°C and 2 h at room temperature. Primary antibody details are as follows: SERCA2a (1:5000, rabbit, A010-20, Badrilla), SERCA1 (1:1000, mouse, CaF2-5D2, DHSB), CSQ1&2 (1:1000, mouse, MA3-913, Thermofisher), CSQ2 (1:1000, rabbit, ab3516, Abcam), ACTN3 (1:0000, rabbit, ab68204, Abcam), SLN (1:1000, rabbit, ABT13, Merck Millipore), MyHC II (1:200, mouse IgG,A4.74, DHSB), MyHC I (1:200, Mouse, IgM, A4.840, DSHB). and actin (1:1000 rabbit, ab1801, Abcam). All antibodies were diluted in LI-COR blocking buffer in PBS (LI-COR Biosciences) 1:1 v/v with 1 × TBST. After incubation in primary antibody, membranes were washed in 1 × TBST, incubated in secondary antibody (1:20,000, IRDye 680-conjugated donkey anti-mouse IgG and IRDye 800-conjugated donkey anti-rabbit IgG (926–68,072, 926–32,213), LI-COR Biosciences) and immunoreactive bands were visualized using infrared fluorescence (IR-Odyssey scanner, LI-COR Biosciences). Band density was analyzed using Image Studio Lite v 5.2 (LI-COR Biosciences). During data analysis, the density of each sample for a given protein was expressed relative to the calibration curve and then normalized to the total protein of each respective lane. SLN in homogenate was normalized against actin, which was not different between RR and XX individuals. Single fiber SLN was normalized against Coomassie stain. The same calibration curve was used across all gels and data are expressed relative to the average of the RR subjects on each gel, which was set to 1.0.

#### Myosin heavy chain composition

MyHC composition was determined by electrophoresis using a protocol adapted from Mizunoya *et al.* ^42^. The whole muscle homogenates were diluted 2.5 times in MilliQ H_2_O and then in 2 × sample buffer containing 100 mM DTT, 4% w/v SDS, 0.16 M Tris-HCl (pH 6.8), 43% v/v glycerol and 0.2% bromophenol blue. 10 μl of each sample was loaded on a separating gel consisting of 100 mM glycine, 35% v/v glycerol, 200 mM Tris-HCl (pH 8.8), 0.35% w/v SDS, 8.5% w/v acrylamide-*N,N’*-methlyenebisacrylamide (99:1), 0.1% w/v ammonium persulfate and 0.05% v/v *N,N,N’,N’*-tetramethylethylenediamine. The stacking gel consisted of 10% v/v glycerol, 70 mM Tris-HCl (pH 6.8), 4 mM EDTA, 0.34% w/v SDS, 4% w/v acrylamide-*N,N’*-methlyenebisacrylamide (99:1), 0.1% w/v ammonium persulfate and 0.05% v/v *N,N,N’,N’*-tetramethylethylenediamine. After adding lower (0.05 M Tris Base, 75 mM glycine, 0.05% w/v SDS) and upper running buffer (6 × concentrated lower running buffer with 0.12% v/v 2-mercaptoethanol), electrophoresis was run at 4 °C for 40 min at 10 mA and then for 22 h and 20 min at 140 V. After electrophoresis, gels were stained with the SilverXpress Silver Staining Kit (Invitrogen) according to the manufacturer’s instruction. Bands were analyzed using ImageJ software.

#### Immunoprecipitation

For IP, 1 μg anti-RyR1 (ab2868, Abcam) antibody was bound to 12 μL G-protein Dynal® magnetic beads (10007D, Life Technologies) following the manufacturer’s instructions. The lysates were centrifuged at 700 g and 4 °C for 10 min and the protein concentration of the supernatant was determined with the Bio-Rad Protein Assay (#500-0006). The samples were then diluted to 0.5 μg protein per μl and 400 μl of each sample was added to the antibody-bead complex. Samples were incubated overnight at 4 ºC under gentle rotation. After incubation, samples gently washed four times with HEPES buffer. Samples were placed on a magnet rack for removal of buffer. To remove separate peptides from the beads, 50 μl Laemmli buffer (Bio-Rad) with 5% 2-mercaptoethanol were added and samples were heated for 5 min at 95 °C. To remove the beads from the solution, tubes were placed on a magnet and the solution was transferred to fresh tubes. Samples (10 μg / well) were loaded on precast 4-12% Bis-Tris gels (NuPAGE, Invitrogen) and run for 1 h at 150 V. Proteins were transferred to PVDF membranes for 3 h on ice. After blocking in blocking buffer (LI-COR) and TBS-T for 1 h, membranes were incubated overnight with anti-RyR1 (1:5000, mouse, ab2868, Abcam) and anti-FKBP12 (1:1000 rabbit, ab2918, Abcam) antibodies. After washing with TBS-T, membranes were incubated in secondary antibody (1:20,000, IRDye 680-conjugated donkey anti-mouse IgG and IRDye 800-conjugated donkey anti-rabbit IgG (926–68,072, 926–32,213, LI-COR Biosciences) for 1h at room temperature. Membranes were washed three times with TBS-T and bands were visualized using an infrared fluorescence scanner (IR-Odyssey, LI-COR Biosciences). Band densities were analyzed with Image Studio Lite v 5.2 software (LI-COR). Data are expressed as ratios of FKBP12/RyR1 relative to the group mean of the RR group, which was set to 1.0.

### Human Proteomics Analysis

#### Protein extraction and solubilization

Vastus lateralis muscle biopsies were homogenized in 16 μg/ml PBS and briefly centrifuged. After discarding the supernatant, 200 μl lysis buffer (8M Urea, 1% SDS, 50 mM Tris pH 8.5, Roche protease and phosphatase inhibitor) were added to the pellet and samples were vortexed and sonicated on ice. Cell lysates were then centrifuged for 10 min at 4 °C and 15,000 rpm and cleared lysates were transferred to new tubes. The extracted proteins were precipitated with chilled acetone (1:4 vol) at −20 °C overnight and then centrifuged for 20 min at 14,000 g at 4 °C. The protein pellets were dissolved in 40 μl of 8 M urea and a 3 μl aliquot of each sample was diluted 10-fold to BCA-assay. A volume equivalent of 25 μg of proteins from each sample were adjusted to 43 μl with water and 5 μl of 1 M ammonium bicarbonate (AmBic) was added.

#### In solution digestion and TMT-labeling

Proteins were reduced with adding 1.5 μl of 200 mM dithiothreitol (DTT, Sigma) in 500 mM AmBic and incubated at 37 °C for 1 h with shaking at 400 rpm. Alkylation was performed with adding 1.5 μl of 66 mM iodoacetamide (Sigma) in 500 mM AmBic at room temperature for 30 min with shaking at 400 rpm. Thereafter 1 μg of sequencing grade modified Trypsin (Promega) was added to each sample (1:33 trypsin:protein) and incubated for 16 h at 37 °C. The digestion was stopped by adding of formic acid at final concentration of 5% and incubating the solution for 20 min at 37 °C. Then the samples were cleaned on a C18 Hypersep plate (Thermo Scientific), dried using a Speedvac and re-suspended in 70 μl of 50 mM triethylammonium bicarbonate (TEAB) buffer and 30 μl of TMT-10plex (Thermo Scientific) reagent was added in dry acetonitrile (ACN) following incubation for 2 h at room temperature with shaking at 550 rpm. Labeling reaction was quenched with 11 μl of 5% hydroxylamine. Labeled samples were combined and dried on Speedvac. Following a cleaning on StageTip C18 20 μl of the combined samples were dissolved in 0.1% formic acid and 2% ACN.

#### PRLC-MS/MS analysis

Chromatographic separation of peptides was achieved using a 50 cm C18 Easy-C18 column (Thermo Scientific) connected to nanoLC-1000 system (Thermo Scientific). Approximately 1.3 μg peptides were loaded onto the column in a volume of 2 μl and then eluted at a 300 nl / min flow rate for 180 min at a linear gradient from 4% to 26% ACN in 0.1% formic acid. Orbitrap Q Exactive plus mass spectrometer (Thermo Scientific) analyzed the eluted peptides that were ionized with electrospray ionization. The survey MS spectrum was acquired at the resolution of 140,000 in the range of m/z 350-1600. MS/MS data were obtained with a higher-energy collisional dissociation (HCD) for ions with charge z=2-3 at a resolution of 70,000 using m/z 2 isolation width.

#### Proteomics data analysis

Data was analyzed on Proteome Discoverer v2.2 (Thermo Scientific) identifying protein in SwissProt database and the extracted abundances were further evaluated using an in-house developed R algorithm calculating fold changes and *P*-values. Data were visualized using Qlucore Omics Explorer (Lund, Sweden).

### RNA-sequencing in BAT of Actn3 KO and WT mice

Brown adipose tissue (BAT) was collected from *Actn3* WT and KO mice after 5 h of core body temperature analyses across the three treatment groups (Thermoneutral, TN, Room-temperature, RT, and cold exposed, CE). A total of 18 WT and 16 KO mice underwent RNA sequencing using the Illumina HiSeq 2500 platform as per manufacturer instructions. Raw read data was processed using the Illumina BaseSpace RNA Express application (Illumina Inc. 2016). Briefly, sequencing reads were aligned using STAR ultrafast RNA seq aligner ^43^ in the SAM file format ^44^, then counted using HTSeq ^45^. The resulting genewise count data was analyzed using the R (3.6.0) statistical programming language (R Core Team 2018). Modelling of differential expression was conducted using the voom precision weights approach ^46^ in the limma (3.40.6) ^47^ package from the Bioconductor (3.9) project.

### Statistics

Statistical analyses were performed using GraphPad Prism 8. Unpaired t-tests were used to assess statistical significance between the RR and XX group. Mantel-COX log-rank test was used for comparisons of the effect of cold-water immersion (baseline to end-point). Data are presented as mean ± SEM and *P* ≤ 0.05 was considered statistically significant.

## SUPPLEMENTARY RESULTS

Table 1 shows physical characteristics and baseline temperatures of individuals in the RR and XX groups. RR subjects were slightly younger than XX subjects, whereas no statistically significant differences were observed for the other measured physical properties or baseline temperatures.

**Supplementary Table 1.** Physical characteristics and baseline temperatures of the individuals in the RR and XX groups.

|  | ACTN3 genotype groups |  |
| --- | --- | --- |
|  | RR (n=27) | XX (n=15) |
| Age, yrs | 22 ± 1 | 28 ± 2* |
| Height, cm | 181 ± 1 | 183 ± 2 |
| Body weight, kg | 77.2 ± 2.5 | 81.2 ± 3.5 |
| Body surface area, m<sup>2</sup> | 1.96 ± 0.03 | 2.01 ± 0.05 |
| Mean skinfold thickness, mm | 12.1 ± 0.9 | 14.5 ± 1.3 |
| Body mass index, kg·m<sup>-2</sup> | 23.6 ± 0.6 | 24.3 ± 0.7 |
| Body fat, % | 15.7 ± 1.0 | 17.9 ± 1.1 |
| Rectal T, °C | 36.9 ± 0.1 | 36.8 ± 0.1 |
| Calf muscle T, °C | 35.6 ± 0.1 | 35.2 ± 0.2 |
| Skin T, °C | 32.1 ± 0.1 | 31.8 ± 0.2 |

Values are mean ± SEM. \* $P < 0.05$ in unpaired t-test.

**Supplementary Figure 1.**
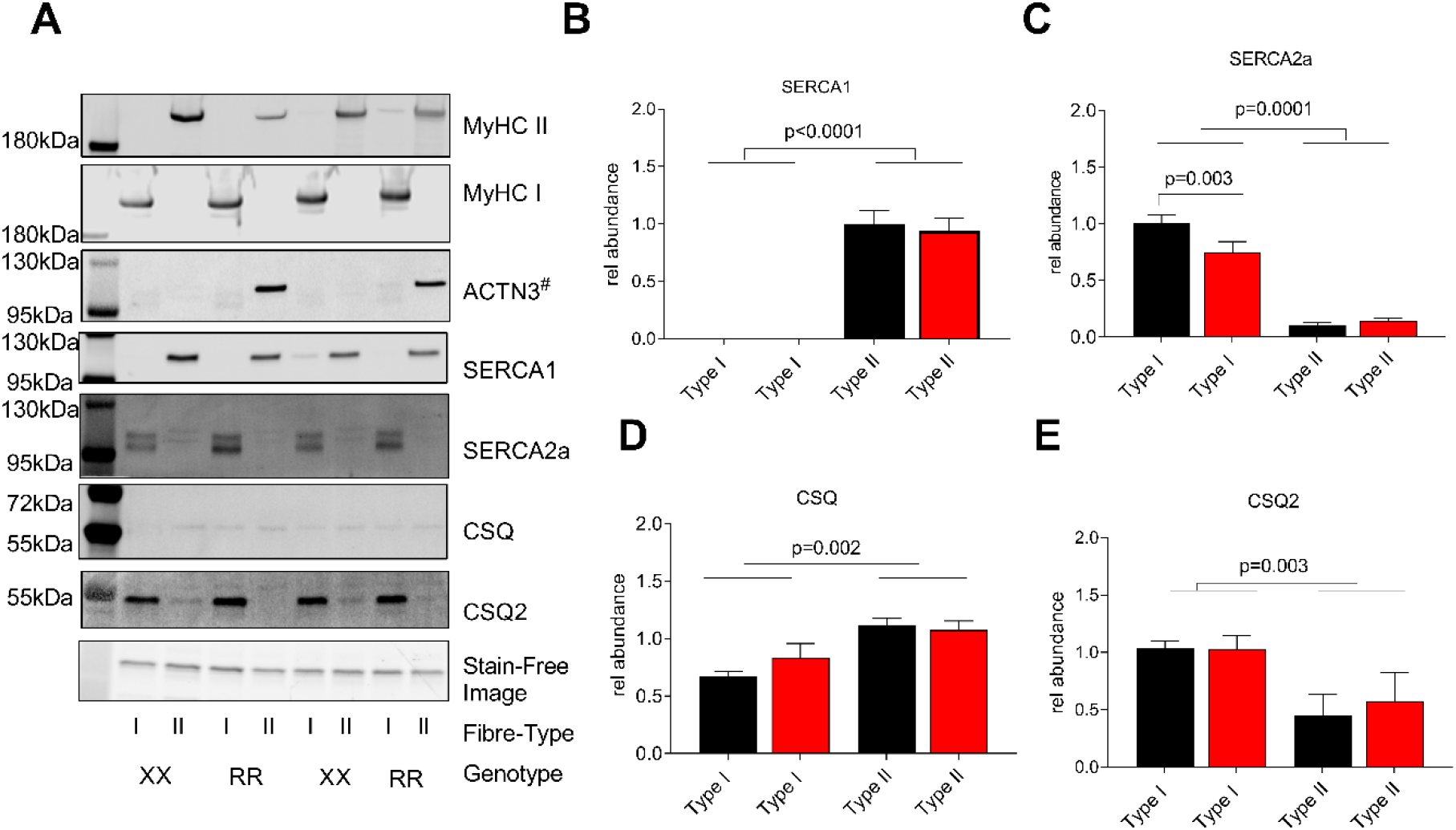
Representative blots (**A**) and summary data of pooled single fiber expression of SERCA1 (**B**), SERCA2a (**C**), CSQ (**D**), and CSQ2 (**E**). Fibers were identified as either Type I (MyHC I) or Type II (MyHC II). ACTN3 has a similar molecular weight as both SERCA1 and 2a, hence it was not possible to probe for ACTN3 on the same membrane as SERCA1 and 2 and an extra gel was run for ACTN3 (marked with #). Stain free images show the actin prior to transfer, which was used as a loading control. For each single fiber pool, proteins were normalized against their own calibration curve (~5-40μg wet weight protein) and total protein and expressed relative to the mean of the RR fibers, which was set to 1; proteins mainly expressed in Type I fibers were expressed relative to the mean of the RR Type I fibers and vice versa for proteins mainly expressed in Type II fibers. Data are presented as mean ± SEM. Differences between RR and XX were determined by one-way ANOVA with Tukey’s *post hoc* test.

**Supplementary Figure 2.**
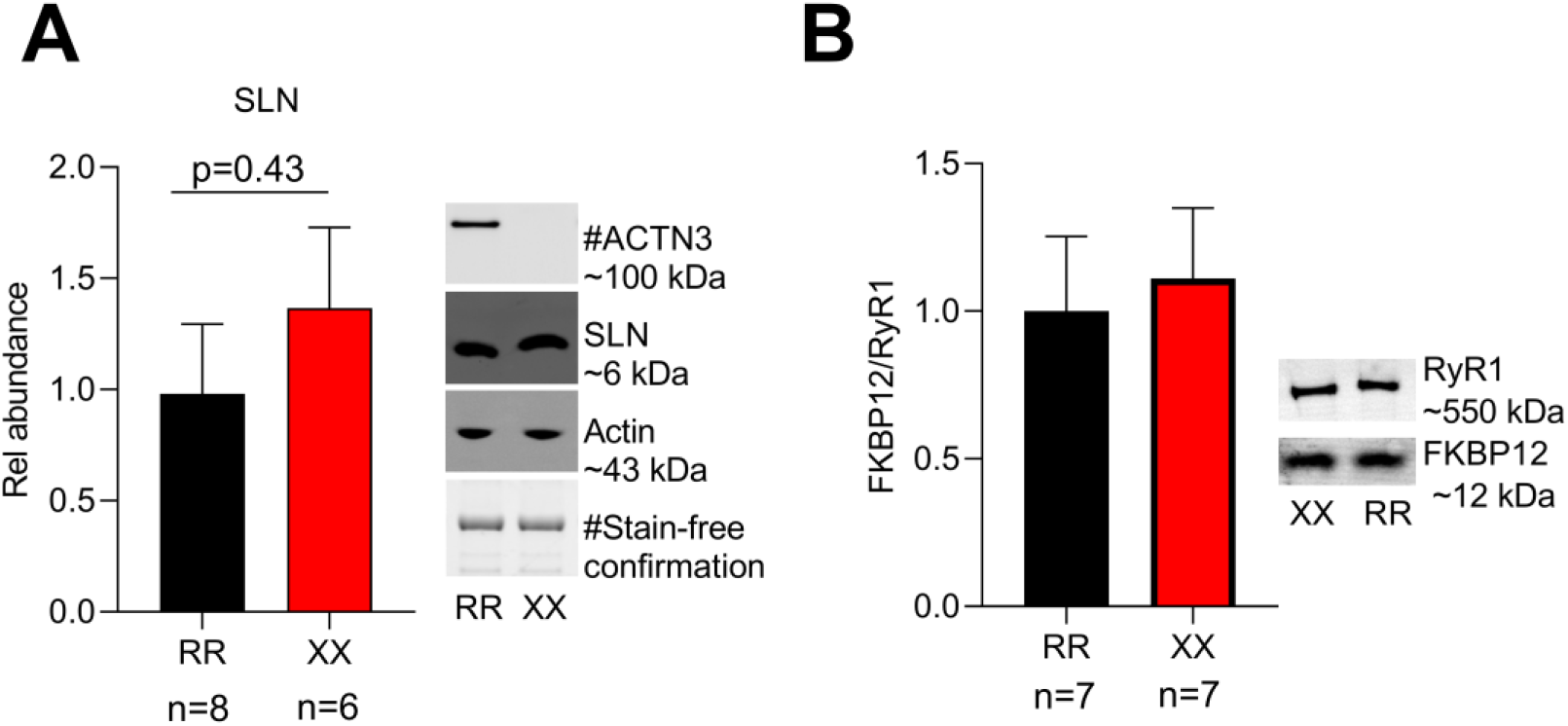
Summary data (mean ± SEM) and representative whole muscle homogenate western blots of SLN (**A**) and FKBP12 associated with RyR1 (**B**). Band intensities in **A** were normalized to their respective actin stain-free loading controls; # in **A** indicates a separate gel from that of SLN (qualitatively similar results obtained with actin on same gel used as loading control). Data are expressed relative to the mean value of the RR group, which was set to 1.0. No statistical difference (*P* > 0.05) in SLN expression or FKBP12 associated with RyR1 between RR and XX individuals were observed with unpaired t-test.

**Supplementary Figure 3.**
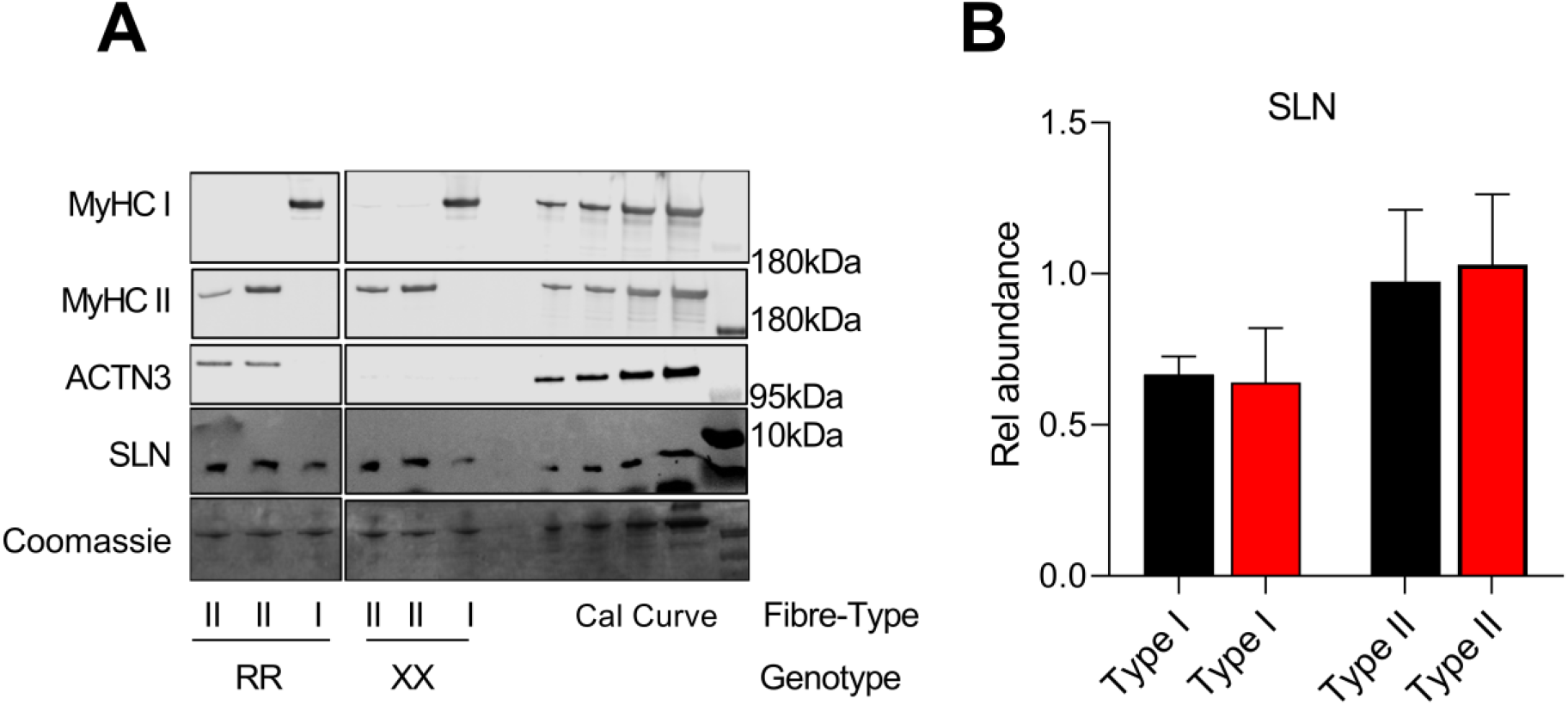
**A**) Representative blots for analysis of SLN protein expression in pooled single fibers. SLN was run on a 16.5% Tris Glycine Gel and Coomassie stained after transfer for loading control. The two MyHC isoforms and ACTN3 were run on two different gels, loaded identical to those described for SLN. Fibers are identified as either Type I (MyHC I), Type II (MyHC II). **B**) For each single fiber pool, SLN was normalized against its own calibration curve and protein content and expressed relative to the mean of the RR MyHC II, which was set to 1. Data are shown as mean ± SEM.

